## Supplementary material for "Retrieving High-Resolution Information from Disordered 2D Crystals by Single Particle Cryo-EM": Suppl. Material

**Running title:** Conformational heterogeneity in 2D crystals

### Supplementary Figures

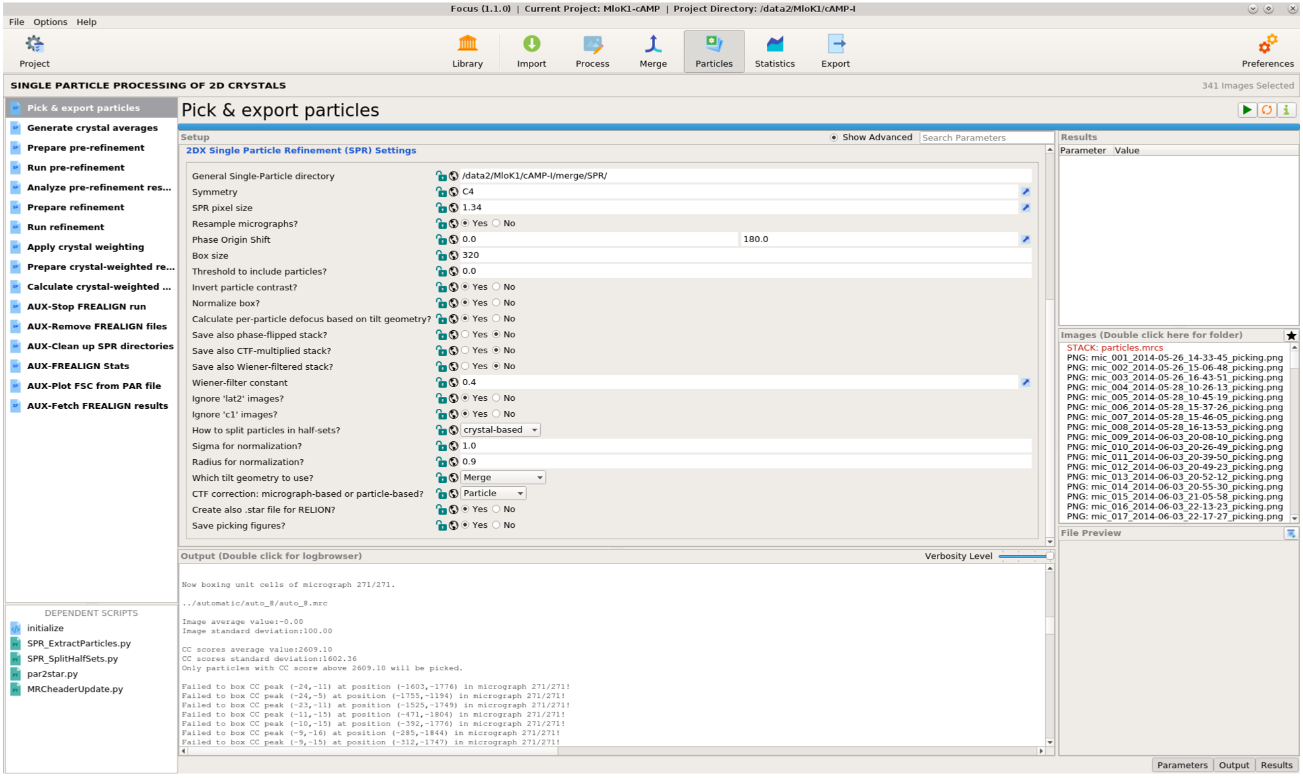


**Supplementary Figure 1. New graphical user interface in FOCUS created to export a 2D electron crystallography dataset for single particle analysis.** The full pipeline for refinement of a single map based on FREALIGN can be executed following the sequence of scripts shown in the panel on the left. Alternatively, all scripts can be called from the command line.

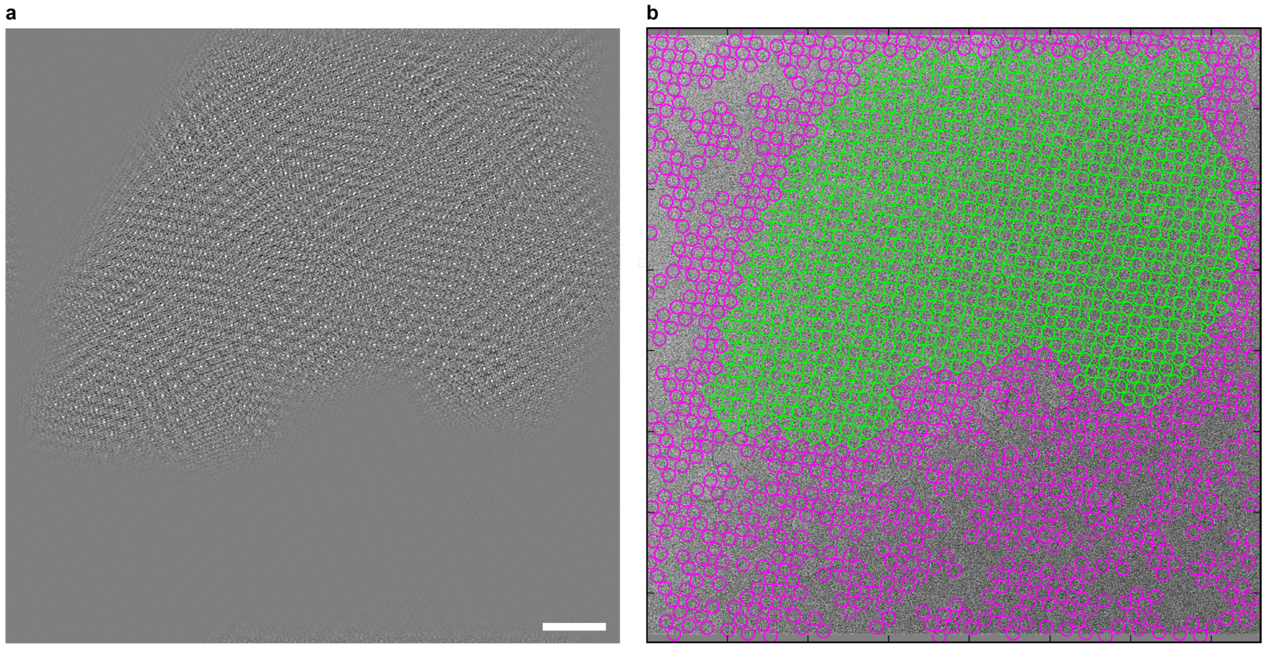


**Supplementary Figure 2. Particle picking from 2D crystals.** The center of each particle (a windowed patch of the 2D crystal) corresponds to the center of each unit cell determined by the classical unbending algorithm[1, 2], optionally with an additional phase shift applied to translate the center of a protein to the center of the window. a) The cross-correlation (CC) profile of a 2D crystal of Mlok1 obtained by the classical unbending procedure[5]. The CC peaks indicate putative positions of unit cells. b) After application of a threshold to the values of the CC peaks to facilitate particle selection: green, particles picked (above threshold); magenta, particles ignored because they are probably bad or false positives (below threshold). Scale bar: 500 Å.

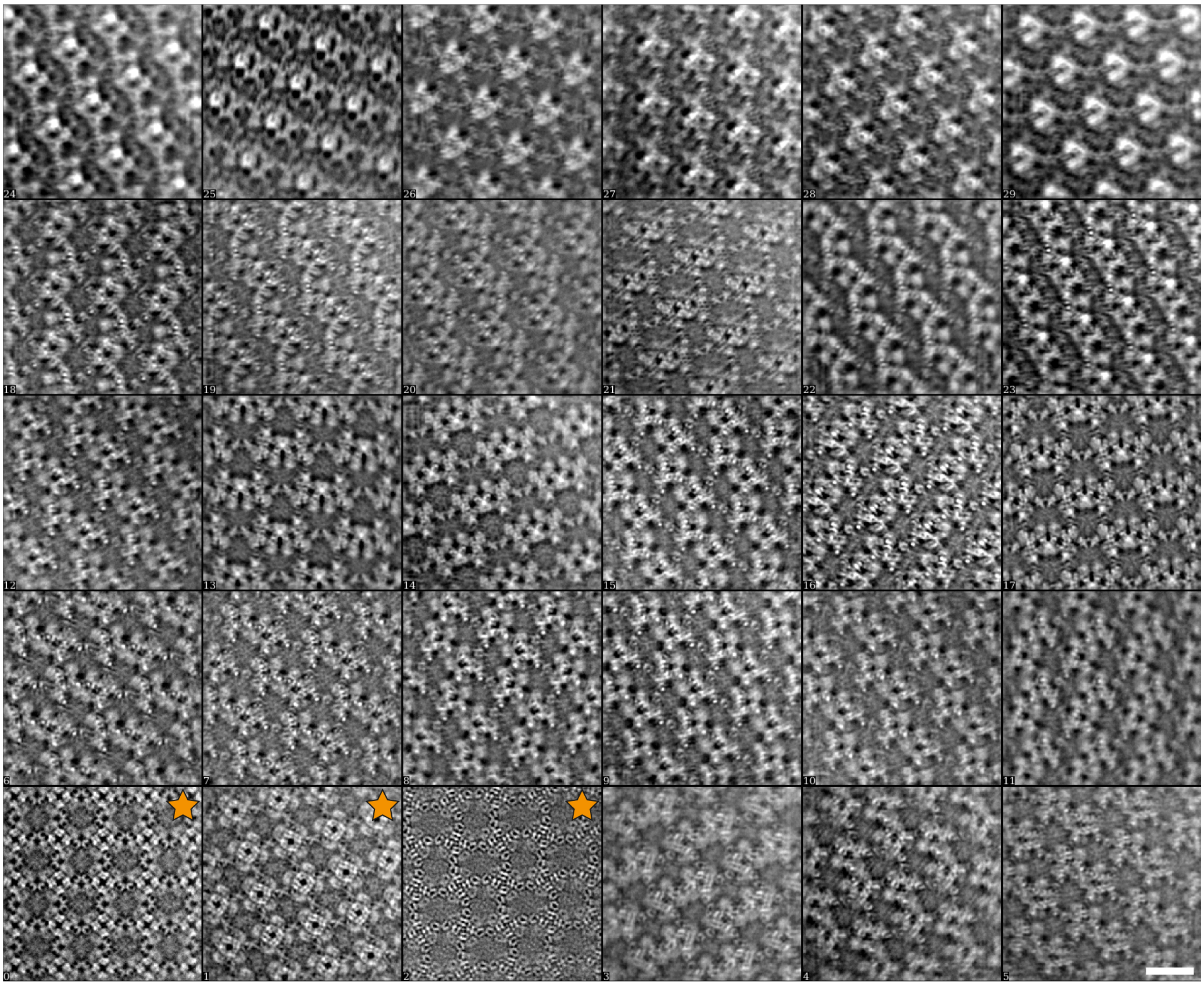

**Supplementary Figure 3. Randomly selected crystal averages from the MloK1 dataset.** The particles contain one MloK1 tetramer in the center and at least eight neighboring complete tetramers, depending on the crystal tilt angle. Images of non-tilted samples (tilt angle < 5°) are marked with a star. Scale bar: 100 Å.

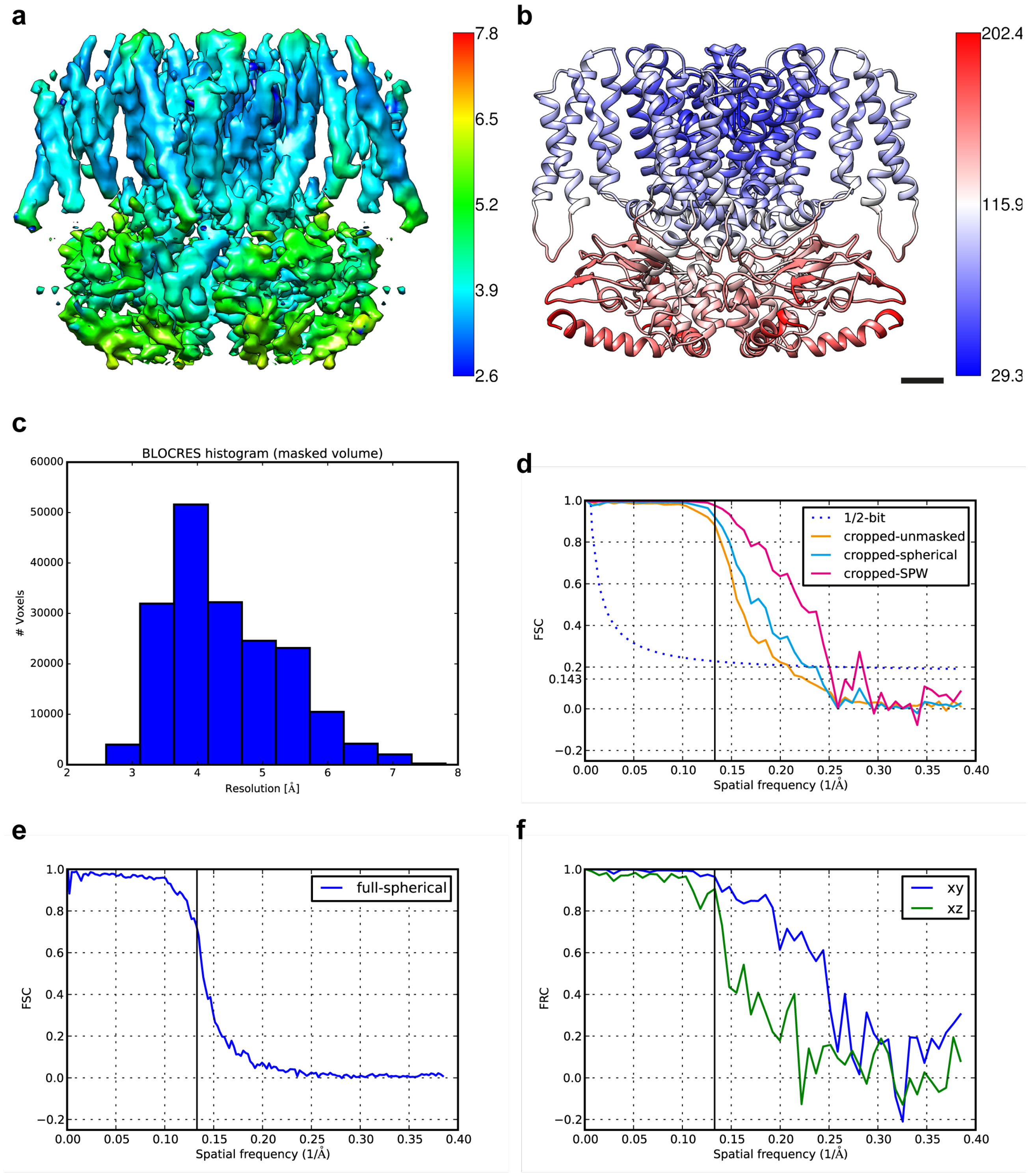


**Supplementary Figure 4. Resolution of the consensus map.** a) Local resolution map according to Blocres using a kernel of 20 cubic voxels and an FSC threshold of 0.143 with colorbar units in Å. b) Average B-factor per residue of the refined MloK1 atomic model with colorbar units in Å$^2$. c) Histogram of voxel-assigned resolutions according to Blocres as in a). d) FSC of the central area cropped for postprocessing and analysis of the central MloK1 tetramer, with a box size of 104 cubic voxels. The solid lines correspond respectively to: orange, FSC between the unmasked cropped half-volumes; cyan, FSC between the cropped half-volumes after applying a soft-edged spherical mask of radius 54.6 Å; magenta, using the same soft-edged spherical mask, but after adjusting the masked FSC for the relative volumes of the molecule (MW=160 kDa) and the mask according to the Single Particle Wiener (SPW) Filter[3] (Fpart/Fmask = 0.288). For resolution assessment, both the 0.143 cutoff and the ½-bit criterion curve[4] (blue dotted line) are shown (calculated for 4-fold symmetric particle with longest dimension of 100 Å). e) FSC between full half-maps at the end of consensus refinement, corresponding to a box size of 320 cubic voxels (i.e., before cropping) and masked by a soft-edged sphere with radius of 192.96 Å. f) FRC between the unmasked, cropped half-volumes as in d) along orthogonal central slices only: blue, along the *xy*-plane; green, along the *xz*-plane (identical to the FRC along the *yz*-plane due to imposition of C4 symmetry, not shown). The solid vertical black line shown in panels d), e) and f) indicates the highest resolution limit used to align the particles in FREALIGN: 7.52 Å.

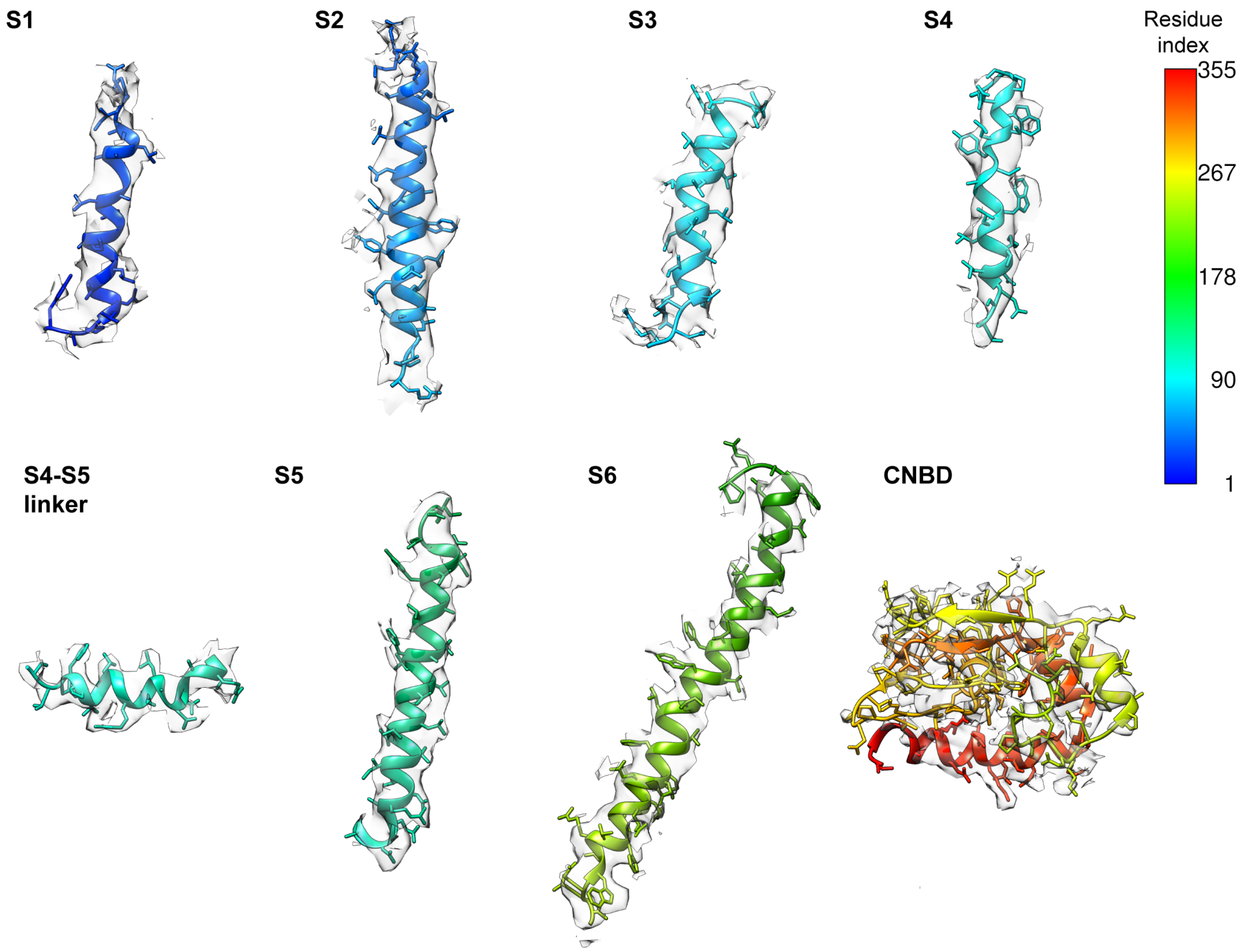


**Supplementary Figure 5. MloK1 model fit to map.** Selected fragments along one chain of the refined MloK1 atomic model shown inside the consensus electron density map. The map was globally sharpened using the *phenix.auto_sharpen* program[5]. A zone of 2.5 Å around the model was used to mask the map in UCSF Chimera[6]. Color bar indicates the residue index from the N- to the C- terminal.

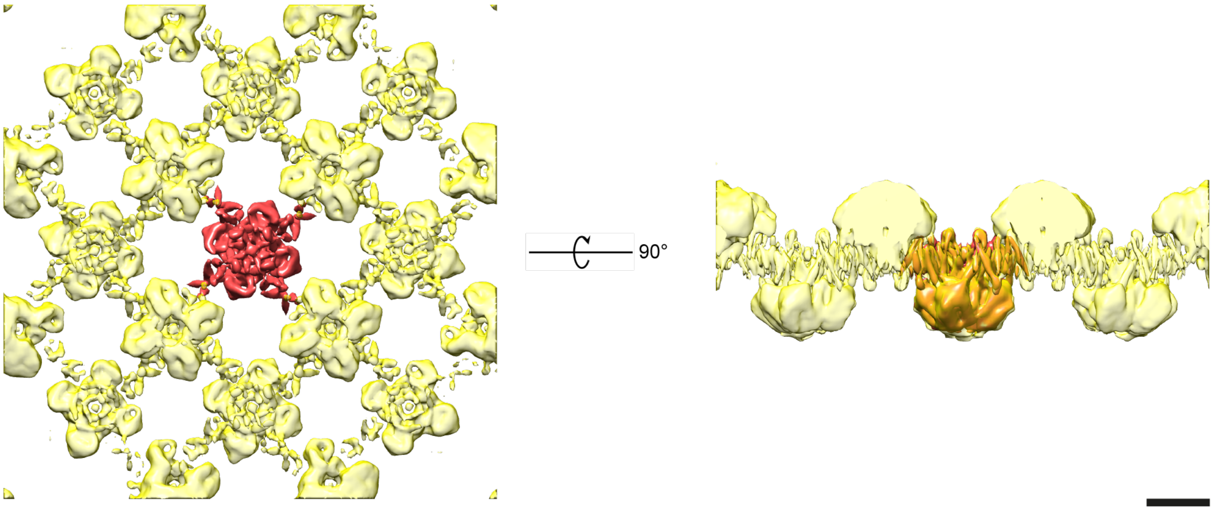


**Supplementary Figure 6. Signal subtraction.** The central MloK1 tetramer, shown in red, was masked using a soft spherical mask. This mask was inverted in order to mask all the neighboring tetramers, shown in transparent yellow, from a fully unmasked reconstruction of the consensus map. This masked map without the central tetramer was subtracted from all experimental particle images using the alignments determined in the consensus refinement, prior to the 3D classification. For this operation, the map and the particles were coarsened by a factor of 2 (*i.e.,* a pixel size of 2.6 Å) by Fourier cropping. Scale bar: 50 Å.

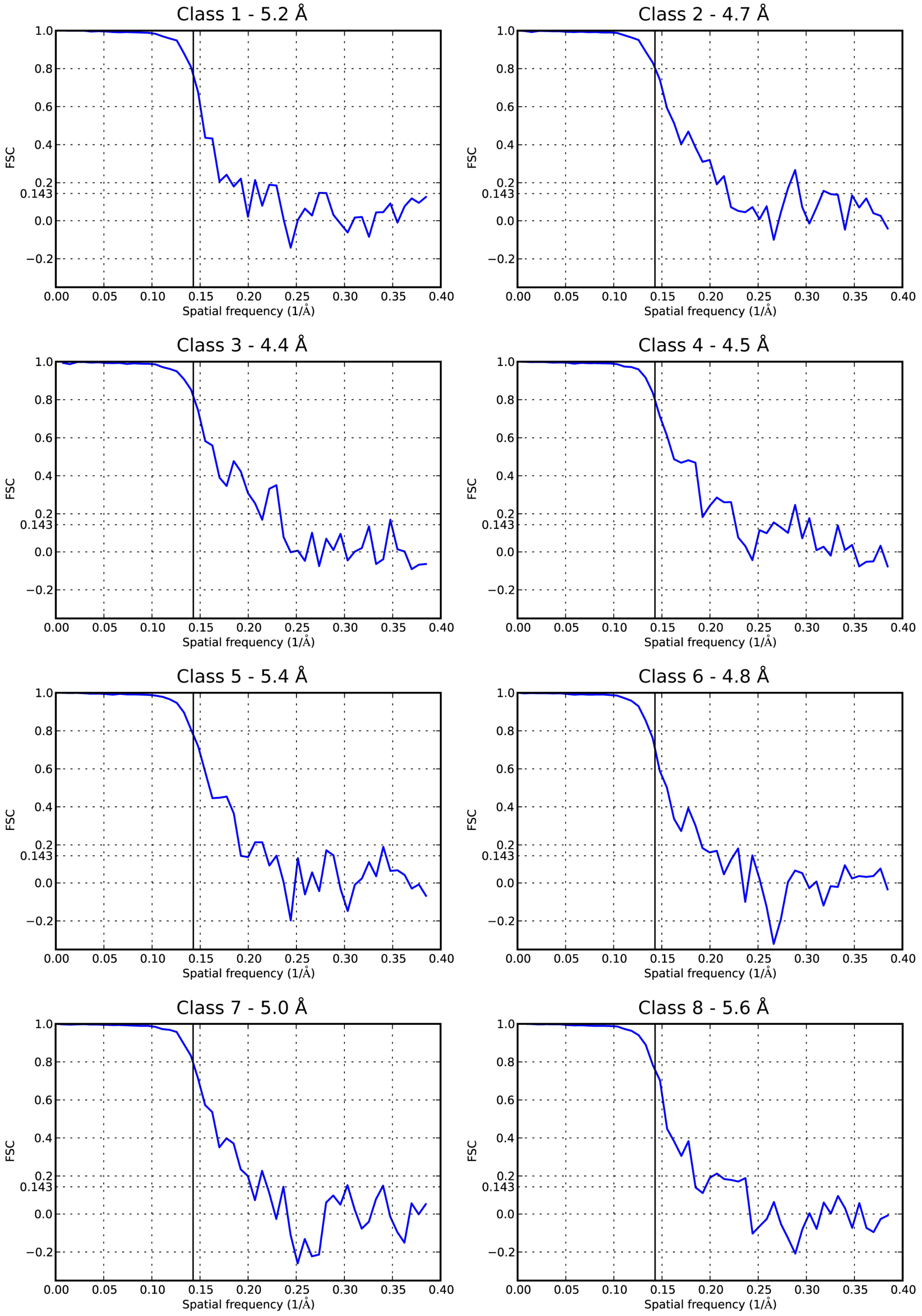
Class 1 - 5.2 Å
Class 2 - 4.7 Å
Class 3 - 4.4 Å
Class 4 - 4.5 Å
Class 5 - 5.4 Å
Class 6 - 4.8 Å
Class 7 - 5.0 Å
Class 8 - 5.6 Å
FSC
1.0
0.8
0.6
0.4
0.2
0.143
0.0
−0.2
Spatial frequency (1/Å)
0.00
0.05
0.10
0.15
0.20
0.25
0.30
0.35
0.40

**Supplementary Figure 7. Resolution of the 3D classes.** FSC curves of the central area cropped for postprocessing and analysis of the central MloK1 tetramer, with a box size of 104 cubic voxels, as performed for the consensus map **(Supplementary Figure 4)**. Likewise, all FSC curves were calculated after masking the half-volumes with a soft-edged spherical mask with a radius of 54.6 Å and adjusted for the relative volumes of the molecules (MW=160 kDa) and the mask according to the Single Particle Wiener filter[7]. The resolution stated next to the class number on the title of each plot corresponds to the 0.143 FSC threshold[7]. The solid vertical black line indicates the resolution limit used to classify the particles in FREALIGN: 7.0 Å. No particle alignment was performed at the stage of 3D classification.

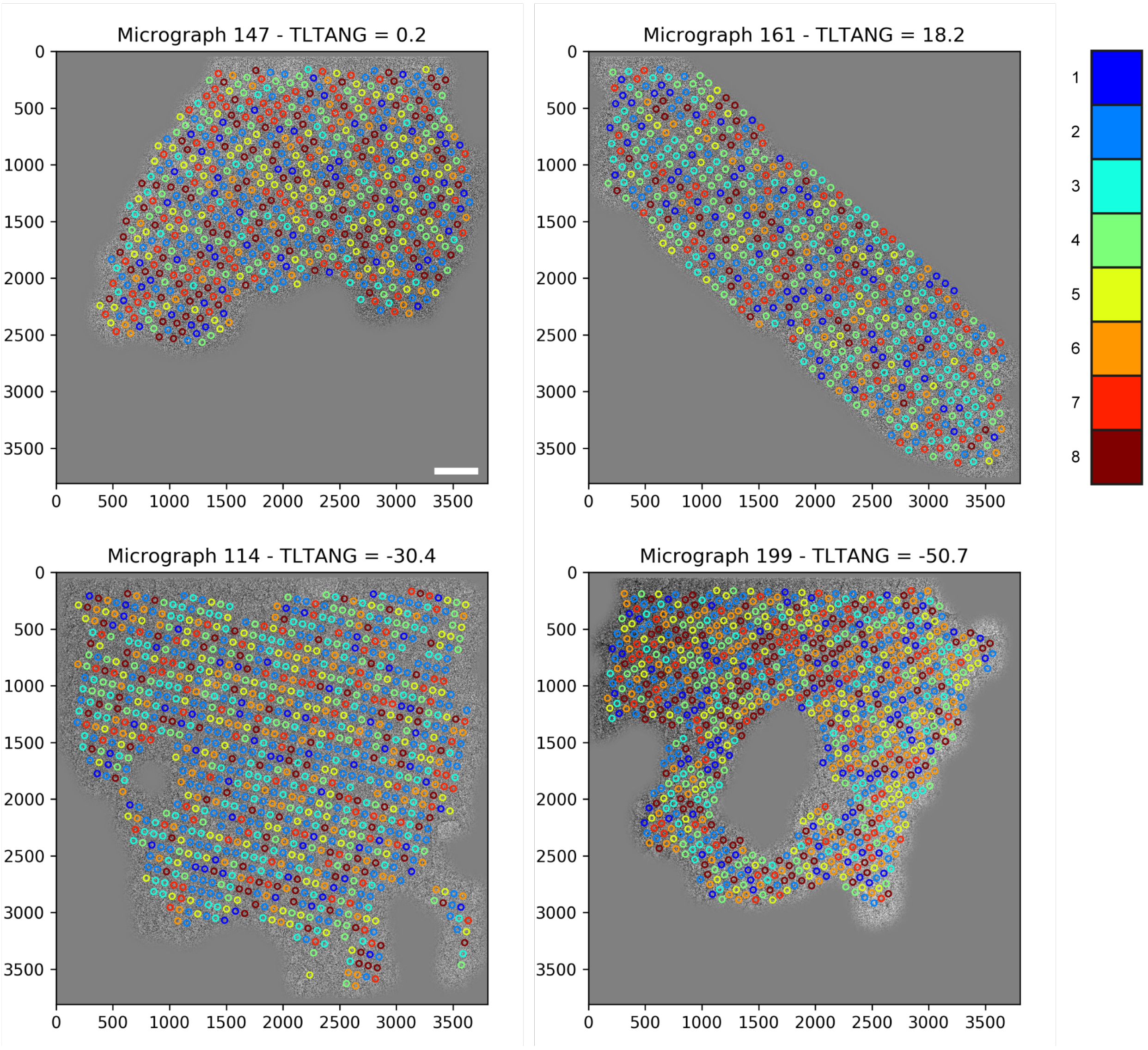


**Supplementary Figure 8. 3D classification results and particle positions.** The locations where particles were picked from the 2D crystals are shown on four representative micrographs of the MloK1 dataset. Each circle around the center of the particle is color coded according to the class with the highest occupancy (maximum likelihood) for that particle. Scale bar is 500 Å.

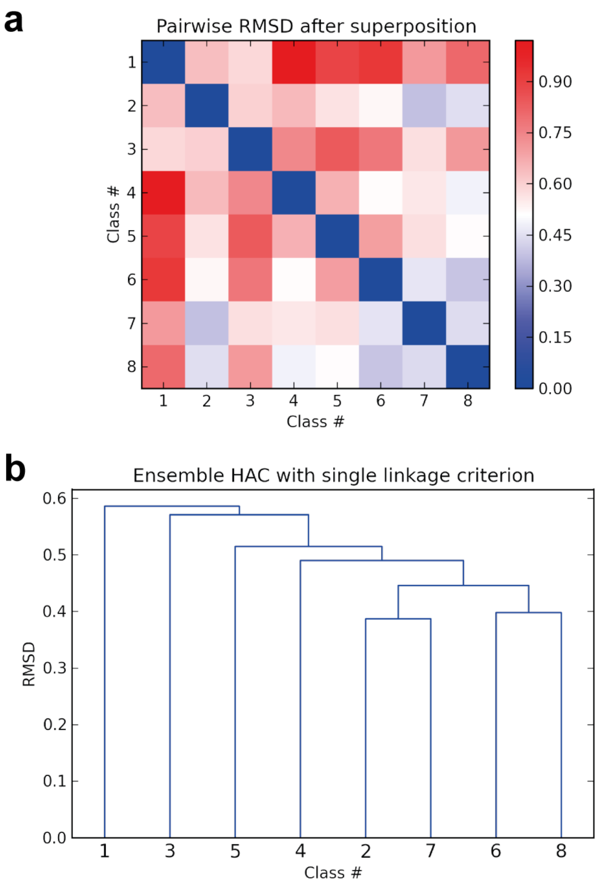


**Supplementary Figure 9. Pairwise similarity within ensemble of atomic models.** a) The pairwise RMSD between the eight atomic models derived from the 3D classes was calculated after superposition of the C-α atoms and plotted as a symmetrical matrix for ease of comparison. b) Based on the RMSD matrix from a), a hierarchical agglomerative clustering (HAC) was calculated using the single linkage criterion[8] and the dendrogram was plotted to indicate similarity relationships within the ensemble. Model #1 is the most different from all other models in the ensemble, followed by model #3; conversely, models #2 and #7 are the most similar to each other, followed by models #6 and #8. In both panels the RMSD values are given in Angstroms. See also **Supplementary Table 2**.

### Supplementary Tables

**Supplementary Table 1. Summary of MloK1 single-particle refinement from 2D crystals.** Values in parentheses refer to the previously published structure[9].

| | |
|---|---|
| **Symmetry applied** | C4 ($P42_12$) |
| **Global resolution** | 4.0 Å (4.5 Å) |
| **Number of micrographs** | 270 (346) |
| **Number of particles** | 231,688 |
| **Pixel size** | 1.3 Å |
| **Box size** | 320 |
| **Clashscore** | 6.30 |
| **Ramachandran outliers** | 0.00 % |
| **Ramachandran favored** | 88.31 % |
| **Rotamer outliers** | 0 |
| **Molprobity score** [10] | 1.94 |
| **EMRinger score** [11] | 0.810 |
| **$CC_{mask}$** [12] | 0.677 |

**Supplementary Table 2. Pairwise RMSD values among members of the model ensemble.** Values computed between all C-α atoms after global superposition, sorted in descending order. See also **Supplementary Figure 9**.

| Model # | Model # | RMSD [Å] |
|---|---|---|
| 1 | 4 | 1.021 |
| 1 | 6 | 0.927 |
| 1 | 5 | 0.892 |
| 3 | 5 | 0.845 |
| 1 | 8 | 0.808 |
| 3 | 6 | 0.781 |
| 3 | 4 | 0.742 |
| 3 | 8 | 0.710 |
| 1 | 7 | 0.708 |
| 5 | 6 | 0.701 |
| 4 | 5 | 0.661 |
| 2 | 4 | 0.643 |
| 1 | 2 | 0.638 |
| 2 | 3 | 0.601 |
| 1 | 3 | 0.586 |
| 3 | 7 | 0.571 |
| 5 | 7 | 0.571 |
| 2 | 5 | 0.563 |
| 4 | 7 | 0.558 |
| 2 | 6 | 0.525 |
| 4 | 6 | 0.518 |
| 5 | 8 | 0.515 |
| 4 | 8 | 0.490 |
| 6 | 7 | 0.461 |
| 2 | 8 | 0.448 |
| 7 | 8 | 0.446 |
| 6 | 8 | 0.398 |
| 2 | 7 | 0.387 |

#### Supplementary Movies

**Supplementary Movie 1.** Blinking of the CNBD. A simple morph from model #4 (“compact” conformation) to model #1 (“extended” conformation) in the ensemble derived from the 3D classes. Each chain is shown with a different ribbon color. Potassium ions are colored purple, and the side chains in the selectivity filter are explicitly shown (residues 175-178). a) Side view; b) CNBD view; c) pore view. Movie generated in UCSF Chimera[6].

**Supplementary Movie 2.** Blinking of the CNBD and tilting of the VSD. A simple morph from model #6 (“compact” conformation) to model #1 (“extended” conformation) in the ensemble derived from the 3D classes. Each chain is shown with a different ribbon color. Potassium ions are colored purple, and the side chains in the selectivity filter are explicitly shown (residues 175-178). a) Side view; b) CNBD view; c) pore view. Movie generated in UCSF Chimera[6].

**Supplementary Movie 3.** Rotation of the CNBD and the selectivity filter with respect to the TMD. A simple morph from model #5 (intermediate “compact” conformation) to model #3 (intermediate “extended” conformation) in the ensemble derived from the 3D classes. Each chain is shown with a different ribbon color. Potassium ions are colored purple, and the side chains in the selectivity filter are explicitly shown (residues 175-178). a) Side view; b) CNBD view; c) pore view. Movie generated in UCSF Chimera[6].

#### Supplementary Notes

##### Alignment restraints in FREALIGN

When processing 2D crystal data with FREALIGN v9.11, it might be necessary to restrain the changes in Euler angles and x,y shifts. We therefore introduced optional restraints to the scoring function being optimized, in the form of Gaussian priors. These restraints have the Gaussian form previously described for x,y shifts and defocus[13] and for helical parameters[14]. Following the notation from Chen *et al.*[13], FREALIGN maximizes a weighted similarity measure $CC_w$ between each particle image $\boldsymbol{X}$ and a projection $\boldsymbol{A}$ of the 3D model:

$$S(\phi;\Theta) = CC_w(\phi;\Theta) + \frac{\sigma^2}{|\boldsymbol{X}||\boldsymbol{A}|}\ln f(\phi;\Theta) \qquad \text{(S1)}$$

where $\phi$ is the set of parameters being optimized, $\Theta$ is a set of parameters governing the restraints imposed on $\phi$ or on a subset of its parameters, $\sigma$ is an estimate of the noise standard deviation equal to $|\boldsymbol{X}-\boldsymbol{A}|/\sqrt{N}$, $N$ is the number of pixels in the image and $f$ is the restraint function. For the x,y shifts of an image, the restraint imposed on refinement cycle $i$ is has the same form as the restraint described in Chen *et al.*[13]:

$$f_{xy}(\phi;\Theta) = \exp\left[-\frac{(x_i - x_{i-1})^2}{2\sigma_x^2} - \frac{(y_i - y_{i-1})^2}{2\sigma_y^2}\right] \qquad \text{(S2)}$$

where $\sigma_x = \sigma_y$ is a parameter defined by the user controlling how strongly restrained the translational change must be. For an Euler angle $\psi$, the restraint is then:

$$f_\psi(\phi;\Theta) = \exp\left[-\frac{(\psi_i - \psi_{i-1})^2}{2\sigma_\psi^2}\right] \qquad \text{(S3)}$$

and analogously for the Euler angles $\theta$ and $\varphi$. For simplicity, we assume $\sigma_\psi = \sigma_\theta = \sigma_\varphi$.

The translational and angular restraints can be specified by their respective keywords in FREALIGN's *mparameters* file as:

### Alignment-restraint parameters

sigma_angles 0.0 ! When greater than 0: Restrains the Euler angles to avoid they change too much in one cycle (STD in degrees).

sigma_shifts 0.0 ! When greater than 0: Restrains the x,y shifts to avoid they change too much in one cycle (STD in Angstrom).

#### Auto-refinement in FREALIGN

We implemented a single-particle auto-refinement algorithm based on FREALIGN v9.11. The algorithm proceeds by evaluating the FSC between the reconstructed half-maps at the end of each refinement cycle at two different thresholds: a “high” threshold (*thresh_fsc_ref*), for example FSC = 0.5, and a lower threshold (*thresh_fsc_eval*), for example FSC = 0.143[7]. For the next refinement cycle, it uses the resolution limit based on *thresh_fsc_ref*. Any FSC improvement beyond this resolution limit, evaluated at *thresh_fsc_eval*, is considered to be unbiased. It is important to remark that these FSC thresholds are arbitrary and not necessarily related to the actual map resolution. If the map does not improve, based on this criterion, then the same refinement cycle can be run again trying different combinations of parameters (PSI, THETA, PHI, SHX, SHY) for refinement (*change_pmask* option), thus changing and/or reducing the dimensionality of the refinement optimization problem. If all parameter combinations have been exhausted and the FSC does not improve from one cycle to the next, refinement is considered to have converged. Even if auto-refinement has converged in the previous step, in some cases, further resolution improvements may still be obtained by modifying other FREALIGN parameters and starting a new auto-refinement procedure from the last cycle of the previous run. We note that a similar auto-refinement strategy was implemented in the *cis*TEM package[15].

The auto-refiner can be tuned by the user via the following keywords in FREALIGN’s *mparameters* file:

### Auto-refinement parameters (only used if calling frealign_run_refine_auto script)

| | |
|---|---|
| thresh_fsc_ref 0.8 | ! Auto-refiner will take the resolution where the FSC crosses this threshold as the limit for the refinement (typically 0.4 to 0.8). |
| thresh_fsc_eval 0.143 | ! Auto-refiner will evaluate map improvement by looking at the resolution where the FSC crosses this threshold. |
| res_min 40.0 | ! Auto-refiner won't ever use a resolution lower than this as the limit for the refinement. |
| ref_stay_away 2.0 | ! Auto-refiner won't ever use a limit for alignment that is less than this value away from the current map's resolution. |
| change_pmask T | ! T or F. Set to T to allow auto-refiner to try different combinations of parameter_mask if necessary. |
| no_theta F | ! T or F. Set to T to keep the tilt angle THETA fixed (always unchanged) in auto-refinement. Useful for 2D crystal data. |

A modified version of FREALIGN v9.11 supporting these features is available at http://www.github.com/C-CINA.
